# Integrity of membrane fluidity and fatty acid environment are necessary for Ebola virus VP40 assembly and release of viral particles

**DOI:** 10.1101/2022.08.24.505195

**Authors:** Souad Amiar, Kristen A. Johnson, Monica L. Husby, Andrea Marzi, Robert V. Stahelin

**Author notes:** **Author Contributions:** S.A., K.A.J. and R.V.S. designed research, S.A. and K.A.J. performed research; M.L.H. and A.M., contributed to experimental design and performed research; S.A., K.A.J., M.L.H., and A.M. conducted data analysis, S.A. and R.V.S. wrote the paper with input from all authors.

## Abstract

Plasma membrane (PM) domains and order phases have been shown to play a key role in the assembly, release, and entry of several lipid-enveloped viruses. In the present study, we provide a mechanistic understanding of the Ebola virus (EBOV) matrix protein VP40 interaction with PM lipids and their effect on VP40 oligomerization, a crucial step for viral assembly and budding. VP40 matrix formation is sufficient to induce changes in the PM fluidity. We demonstrate that the distance between the lipid headgroups, the fatty acid tail saturation and the order between the two leaflets are important factors for the stability of VP40 binding and oligomerization at the PM. Use of FDA-approved drugs (dibucaine, propranolol and trifluoperazine) to fluidize the plasma membrane, destabilizes the viral matrix assembly leading to a reduction in budding efficiency. Lastly, we show that VP40 can tether and cluster lipid vesicles upon protein enrichment at the membrane. This is a new characteristic of the protein, and it opens the door to new avenues of exploration to deepen our understanding of VP40 host interactions and EBOV assembly. Indeed, our findings support a complex assembly mechanism of the EBOV viral matrix that reaches beyond lipid headgroup specificity using ordered PM lipid regions independent of cholesterol.

## Introduction

Ebola virus (EBOV) is a lipid-enveloped filovirus responsible for sporadic outbreaks of an often-fatal hemorrhagic disease in Western and sub-Saharan Africa. During EBOV infection, the dimeric matrix protein known as virion protein 40 (VP40) is the primary viral component responsible for directing the assembly and budding of nascent viral particles from the plasma membrane (PM) (1– 5). This process occurs at the inner leaflet of the PM, where VP40 is ultimately trafficked. Indeed, it has been shown that VP40 binds selectively to phosphatidylserine (PS) and phosphatidylinositol 4,5-biphosphate PI(4,5)P_2_, enriched anionic lipids at the inner PM leaflet (6–8). In addition, VP40 oligomerizes in an anionic lipid dependent-manner to form the viral matrix layer (6, 7, 9). During this process, the organization within the PM has been shown to change including PS exposure to the outer leaflet that may involve host scramblase at budding sites (10, 11), lipid clustering (12) and induced or spontaneous formation of membrane curvatures for nascent virus particle assembly (13, 14). Although the role of PS and PI(4,5)P_2_ headgroups in VP40 binding and assembly at the PM has been well established, physical properties of the PM that are required for efficient virus budding are still unknown.

The lipid composition of each subcellular membrane is unique and participates in correct organelle function (15, 16). This composition is defined by a large heterogeneity of lipid headgroup species, fatty acid tails, sterol levels and phosphatidylinositol species abundance (17-19). In consequence of this heterogeneity, bio-membranes display varying profiles of membrane fluidity. The PM is well known to maintain rigidity by a constant adjustment of its lipid profile to maintain the correct cell integrity. This rigidity is maintained by the abundance of phospholipids with long and saturated fatty acid tails and/or by an enrichment of sterols (cholesterol) and sphingolipids. In addition, the enrichment of cholesterol and sphingolipids of the PM results in the formation of dynamic microdomains often referred to as lipid rafts that give rise to tightly packed and ordered lipid regions (20-22).

Lipid binding of viral proteins may be dependent on lipid microdomain formation and lipid acyl chain composition. For example, HIV-1 Gag was shown to selectively bind liquid ordered (L_o_) membranes enriched with cholesterol (23). HIV-1 Gag binding near to PI(4,5)P_2_ -rich-regions of the PM depends on the myristylation of the matrix domain (MA) of Gag. In model membranes, HIV-1 Gag protein binds liquid disordered (L_d_) membranes in presence of PIP_2_ and can sequester the unsaturated *sn*-2 acyl chain of the PIP_2_ in a hydrophobic cluster of MA-myristate, whilst the saturated *sn*-1 acyl chain remains in the PM (24-26). This sequestration could be the origin of PIP_2_ nanodomains formed at the surface of cells during virus assembly (27). These PIP_2_ nanodomains are stabilized by clusters of cholesterol molecules where the saturated *sn*-1 acyl chain of PIP_2_ can favorably be trapped, to create virus specific PI(4,5)P_2_/cholesterol lipid-enriched membrane environment for more efficient viral assembly (27).

In this study, we report the results of our detailed investigations on the importance of membrane fluidity during EBOV VP40 matrix assembly. We first examined whether VP40 binding to membranes affected membrane fluidity using lipophilic dyes that emit fluorescence depending on the polarity of their environment and the rate of dipolar relaxation of the molecules, the Laurdan and pyrene PA probe (28). Next, we identified the factors involved in the plasma membrane rigidity required for proper VP40 binding, oligomerization, and matrix formation. We showed how an increase in the PM fluidity destabilized the integrity of the viral matrix. By performing an in-depth analysis of VP40-membrane interactions, we demonstrated that VP40 binding to the PM goes beyond the head group specificity and requires an ordered hydrophobic environment within the PM.

## Results

### EBOV VP40 modifies membrane fluidity upon PM binding and oligomerization

First, we aimed to investigate the impact of VP40 binding and assembly on the PM physical properties. To do so, we monitored the general polarization of Laurdan fluorescence in the PM of HEK293 cells expressing EGFP or EGFP-VP40. 24 h post-transfection the cells were stained with Laurdan dye and imaged 30 minutes later. The fluctuations in the polarization of the Laurdan signal correlate with water penetration into the lipid bilayer surface due to the dipolar relaxation effect. These fluctuations indicate the lipid packing and membrane fluidity states of studied membranes (29). In cells expressing EGFP-VP40, the analysis of the Laurdan emission signal showed an increase of fluorescence intensity in the red spectrum (490 nm) at regions of the PM enriched with VP40 (Fig 1A, arrow heads). In contrast, in EGFP expressing cells, the red signal was accumulated in the intracellular membranes. The normalized fluctuations of the generalized polarization factor analysis of the dye at the PM confirms an increase of disordered membranes in EGFP-VP40 expressing cells compared to control cells expressing EGFP (Fig 1B). This indicates a relative overall increase in membrane fluidity (16) due to VP40 PM localization and assembly.

**Figure 1.**
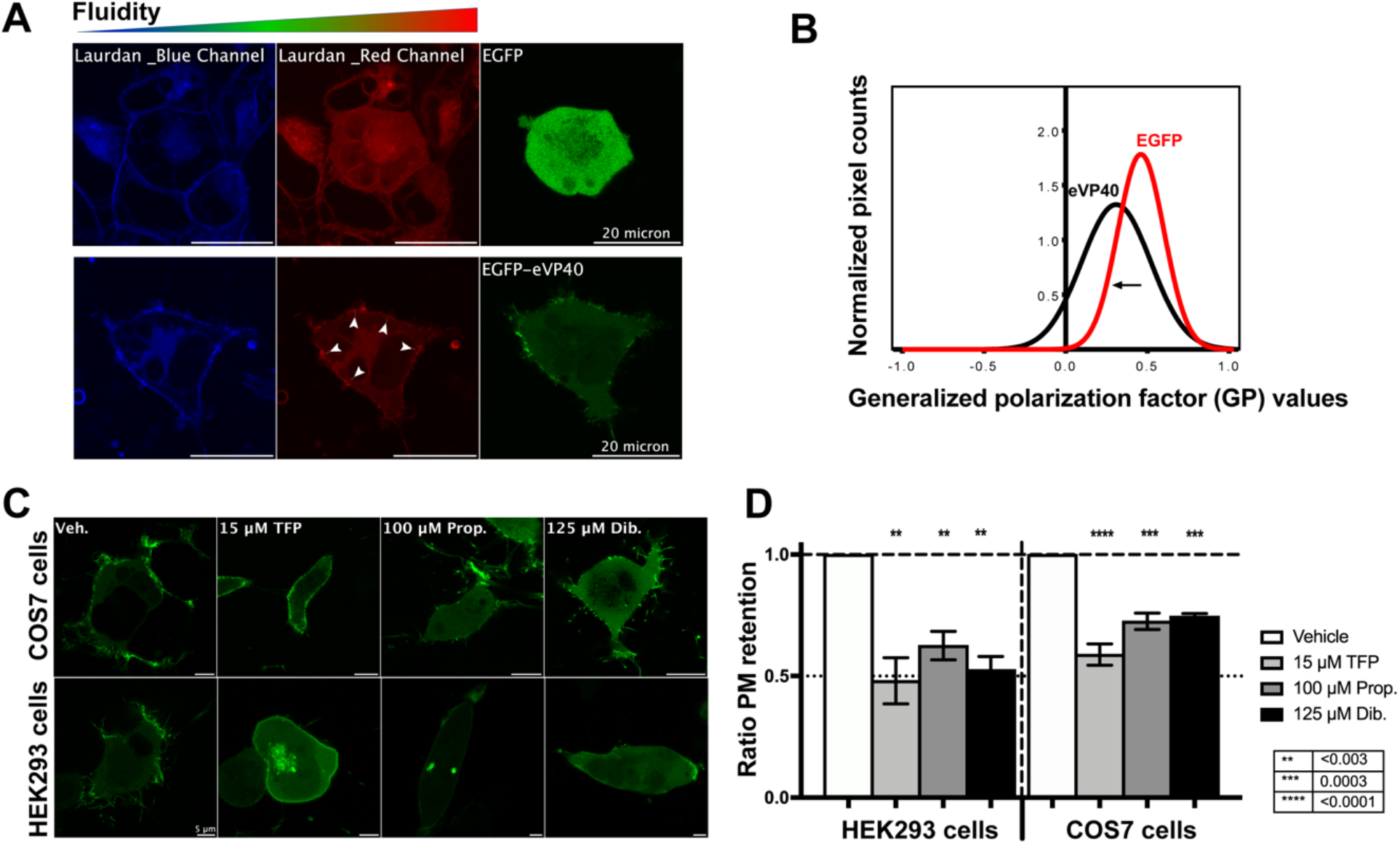
Membrane enrichment of EBOV VP40 alters plasma membrane (PM) fluidity. A). Representative multi-photon confocal images of HEK293 cells expressing EGFP or EGFP-VP40 (green) and treated with Laurdan dye to monitor the change in the PM fluidity in the emission spectrum (blue and red). B) Experimental and fitted normalized generalized polarization distribution curves of Laurdan dye across the PM of HEK293 cells from (A). Generalized polarization values range from −1 (very fluid lipid domains) to +1 (very rigid lipid domains). The fitting procedure was performed using a nonlinear Gaussian curve. C) Representative confocal cell images of live cells expressing EGFP constructs (*green*) treated or not with FDA-approved drugs as indicated for 30 min prior to imaging. D) Ratio of PM retention from (C) quantified by calculating the integrated density of pixels at PM to total pixels within the cell and normalized to vehicle treated cells. The values are reported as mean ± SEM of three independent replicates. One-way ANOVA with multiple comparisons was performed.

PM condensation and cholesterol-dependent microdomain formation have been well documented to be crucial factors for the recruitment of specific proteins important for membrane trafficking and signaling events, such as caveolae formation (20, 30). To determine the influence of increased PM fluidity on VP40 PM association, we used live cell imaging of HEK293 and COS-7 cells expressing EGFP-VP40. PM fluidity was increased using three FDA-approved drugs, dibucaine (Dib.), propranolol (Prop.) and trifluoperazine (TFP) (31-37). All three molecules induced dissociation of VP40 from the PM within 30 min in contrast to vehicle control (30-50%, Fig 1C-E).

VP40 binding to the PM is dependent on the level of PS and PI(4,5)P_2_ at the inner leaflet of the PM (6, 7). LactC2 and PLCδ-PH (a PS and PI(4,5)P_2_ reporter, respectively) were used to check the ability of PS and PI(4,5)P_2_ reporters to undergo typical PM localization (Fig. Supp 1A-B). PM association of these two sensors did not harbor a detectable change of their localization even at higher drug treatment concentrations. These observations indicate the three FDA approved drugs did not impact PS- and/or PI(4,5)P_2_ distribution at the PM and that the changes in VP40 PM association are most likely due to changes in membrane fluidity induces by these drugs.

### VP40 PM association and oligomerization is sensitive to membrane fluidity changes

Here we aimed to understand how membrane fluidity influences VP40 assembly and oligomerization, so we monitored VP40 oligomerization at the PM using two different assays. Number and brightness (N&B) analysis (38) was first performed to assess VP40 oligomerization states at the PM with or without FDA-approved drug treatments. Live cell imaging indicated a rapid reduction in VP40 oligomers, especially larger oligomers (>12mer) in contrast to the vehicle control. Dibucaine also induced an increase of small oligomers (monomer-6mer) at the PM compared to propranolol and TFP (Fig. 2A-B). This analysis indicated that increased membrane fluidity destabilized VP40 oligomerization.

**Figure 2.**
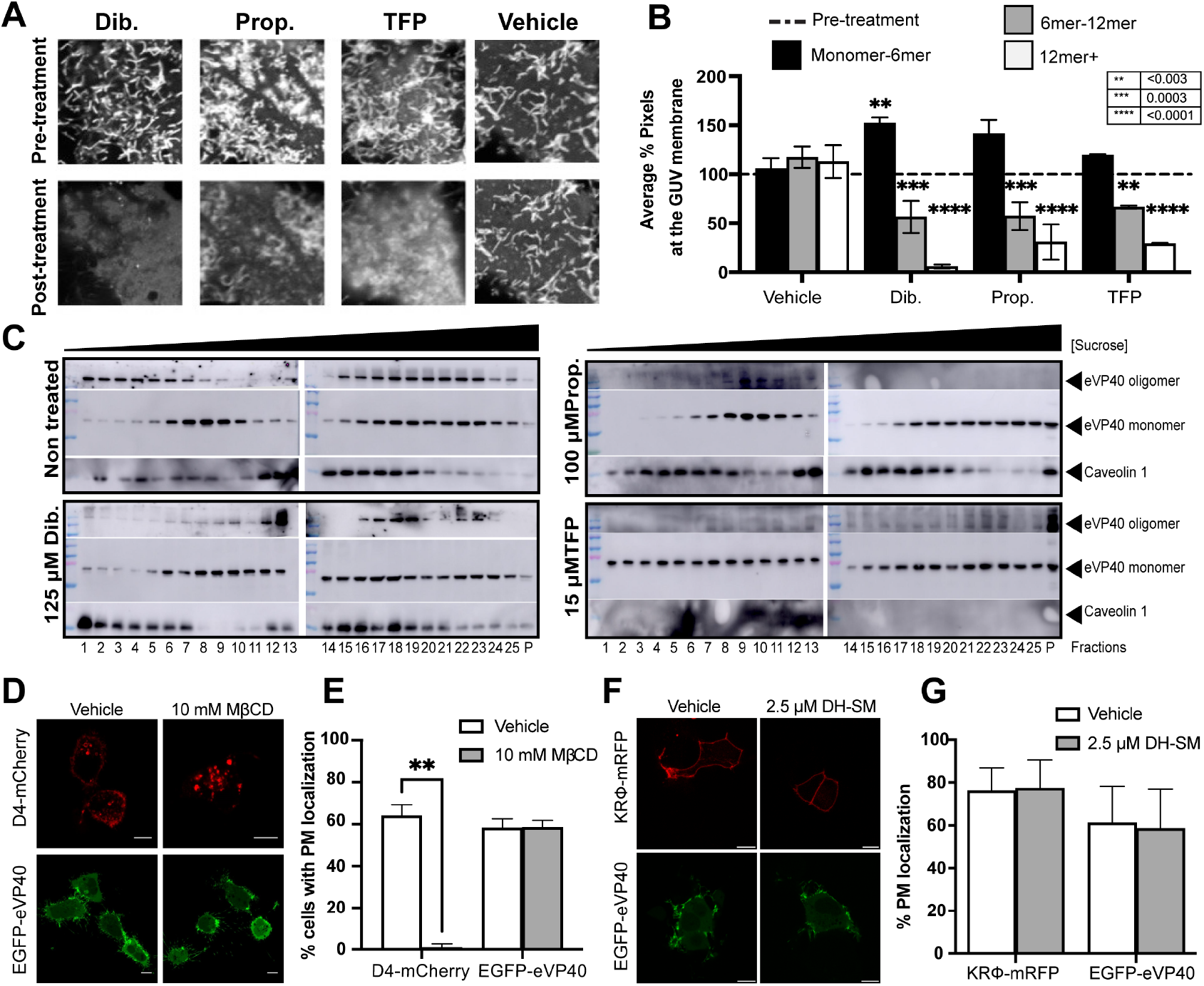
VP40 oligomerization is sensitive to membrane fluidity changes but not to cholesterol extraction. A) Representative images for number and brightness (N&B) analysis of VP40 oligomers pre- and post-FDA-approved drug treatment. B) Distribution of %oligomer populations before and after treatment with FDA approved drugs from (A). Pre-treatment values were normalized to 100% for each oligomer population. The values shown are mean ± SEM of three independent replicates. C) Representative immunoblots of sucrose gradient fractions of detergent-resistant (DRM, including rafts, fractions 1 to 10) and detergent sensitive membranes (DSM, fractions 15-25) isolated from EBOV VP40-expressing HEK293 cells treated for 30 min with FDA-approved drugs as mentioned. D) Representative confocal images of MβCD treatments of cells expressing mCherry-D4 or EGFP-VP40. Scale bars: 5 µm. E) Quantification of total % of cells exhibiting PM localization of the proteins from D), and the values are reported as mean ± SEM of three independent replicates. One-way ANOVA with multiple comparisons was performed. \*\**P*, 0.003. F) Representative confocal images of DH-SM treatments of cells expressing either KRϕ-mRFP or EGFP-VP40. Scale bars: 5 µm. G) Quantification of total % of cells exhibiting PM localization of the proteins from (F), and the values are reported as mean ± SEM of three independent replicates.

A previous study reported that VP40 oligomerization and subcellular localization is associated to membrane microdomains using the detergent-resistant/rafts (DRM) and detergent-sensitive (DSM) membrane fractionation assay (39). To determine the VP40 oligomerization profiles in DSM and DRM fractions upon membrane fluidization, DSM/DRM fractions were prepared by sucrose gradient centrifugation (39–41) from VP40-expressing HEK293 cells treated or not with different drugs for 30 minutes prior to cell lysis. Fractions were then analyzed by Western blotting as previously described (39). In non-drug treated cells, VP40 oligomers were successfully detected in both the DRM (fractions 5-13, along with the DRM marker Caveolin-1 (CAV1)), and DSM fractions (fractions 16-22). Dibucaine and propranolol treatments significantly reduced the amount of VP40 oligomers in DRM and DSM fractions, respectively, while in TFP-treated cells, only monomeric VP40 was detected homogenously across all the fractions (Fig. 2C). Dibucaine and propranolol treatments didn’t disrupt CAV1 subcellular distribution whereas CAV1 was undetectable in TFP-treated cells suggesting that TFP abolished raft and microdomain distribution across the PM (34). This led us to hypothesize that VP40 oligomerization is independent of PM microdomains and that there is likely another factor(s) of PM rigidity for stabilization of the VP40 oligomerization/matrix.

We also investigated if the membrane fluidization effects of these three FDA-approved drugs on VP40 PM dissociation and matrix disassembly were permanent. We combined total internal reflection fluorescence microscopy (TIRFM) with a liquid perfusion system. Herein we treated cells with low to high drug concentrations for 5 min, followed by 8 to 10 min wash steps with calcium-free media. At low concentrations, washing the three drugs from cells resulted in the recovery of VP40 binding and assembly at the PM. In most cases, the re-association of VP40 occurred at the same plasma membrane regions prior to drug treatments (Fig. S1C-E). These data show that the impairment of the PM fluidity is reversible as is VP40 reassociation to the PM.

### EBOV VP40 binds membranes in a cholesterol independent manner

The biophysical properties of the PM rigidity that promote VP40 binding, oligomerization and budding are still unclear. We first depleted cholesterol from the PM using methyl-β-cyclodextrin (MβCD), a widely used reagent to extract cholesterol from PM lipid rafts to form intracellular soluble inclusion complexes (42-45). As shown in Fig. 2D-E, the cholesterol depletion from the PM had no effect on VP40 PM binding, accumulation, and the elongation of virus-like particles (VLP) from the PM. The cholesterol sensor Domain 4 (D4) of Perfringolysin O (46), fluorescently labelled with a mCherry tag, showed a clear delocalization from the PM upon MβCD treatment in all treated cells (Fig. 2D). Lidocaine is another FDA-approved drug that had been demonstrated to increase the membrane fluidity of raft and raft-like domains, decreasing membrane thickness and depleting cholesterol from erythrocyte membranes (47-50). Treating cells expressing VP40 with a high concentration of lidocaine did not affect the protein binding to the PM (Fig. S1A-B). Altogether, the results described here align with our previous observations and confirm our hypothesis that cholesterol and PM cholesterol-rich microdomains are not critical factors in VP40 membrane binding and oligomerization.

Second, we explored the functional effect of increasing the PM rigidity with dihydrosphingomyelin (DH-SM). Previous studies using artificial membranes support that DH-SM:cholesterol-containing liposomes form more condensed liquid ordered (*L*_*o*_) domains than those containing SM (50-55). The cell treatment with DH-SM did not affect the surface potential of the PM as the anionic charge sensor probe KRФ (56) showed a comparable PM localization in vehicle treated cells (Fig. 2F-G). The effect of the DH-SM on the PM rigidity increase had not been reported in cells in previous studies, therefore, we confirmed it in this study where the Laurdan generalized polarization factor showed a shift toward more rigid membranes when cells were treated with 2.5 µM DH-SM (Fig. S2). The PM rigidity increase exhibited no significant effect on VP40 PM localization (Fig. 2F-G). Altogether, our data suggest the membrane rigidity required for VP40 association and stabilization of the matrix formation may be independent of cholesterol-sphingolipid domains.

### Effects of acyl chain saturation on VP40 binding to membranes

We next assessed the role of phospholipid fatty acid tails in VP40 membrane binding. We measured the *in vitro* association of VP40 to membranes of different fluidity, namely liquid disordered (L_d_; DOPS:DOPC:DOTAP), liquid ordered (L_*o*_; DPPS:DPPC:Cholesterol) and solid ordered (S_o_; DPPS:DPPC) membranes. VP40 poorly associated with membranes of L_d_ origin but binding was significantly increased to L_o_ and S_o_ membranes containing PS (Fig 3A). In all cases, more rigid membranes increased VP40 binding whereas cholesterol did not play an important role as S_o_ membranes were devoid of cholesterol, confirming our previous observations from cellular studies. As PI(4,5)P_2_ is an important factor for VP40 membrane binding, oligomerization, and particle formation, we assessed the influence of PI(4,5)P_2_ on VP40 membrane association. Indeed, PI(4,5)P_2_ increased VP40 membrane association in both L_d_ and L_o_ membranes but a clear selectivity for ordered membranes by VP40 was observed (Fig 3B-C). This indicates that acyl chain saturation or membrane rigidity (which can be dependent on acyl chain composition) are important determinants of VP40 membrane association.

**Figure 3.**
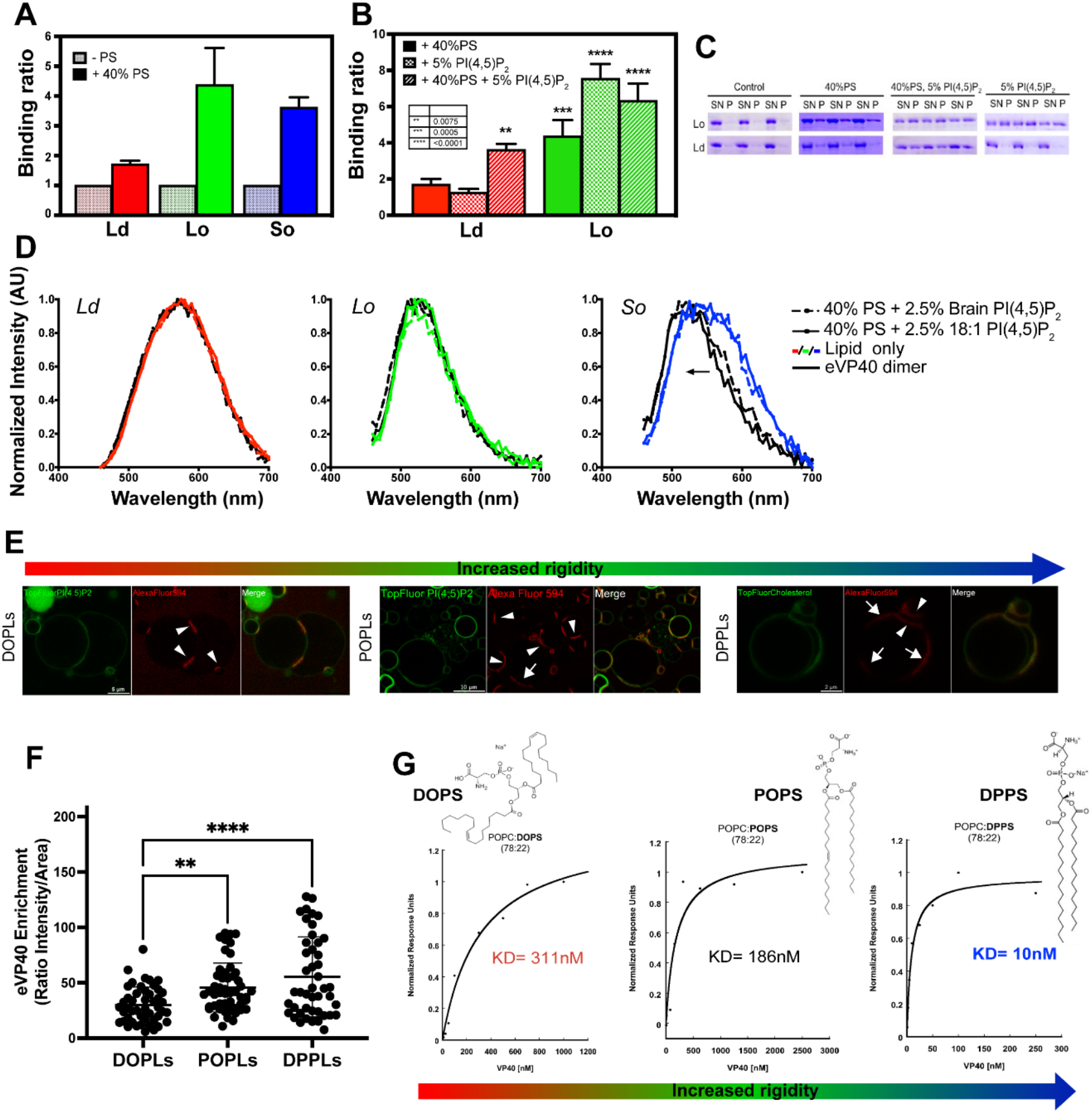
Recombinant EBOV VP40 exhibits significant binding specificity for rigid membranes driven by the saturation level of phosphatidylserine (PS). Lipid pelleting assay analysis shows fold change in EBOV VP40 bound to (A) PS- and/or (B) PI(4,5)P_2_-containing liposomes with different membranes phases: liquid disordered (L_d_), liquid ordered (L_o_) and solid ordered (S_o_) compared to control membranes (without anionic lipids). The values are reported as the standard error of the means and two tailed students T-test has been performed. (C) Representative SDS-PAGE bands of VP40 in the supernatant (SN) and pellet (P) fractions of the lipid pelleting assay. D) Membrane fluidity assay performed on lipid vesicles using a PA pyrene probe (28). Normalized absorption and fluorescence spectra of the PA probe in LUVs of different composition and lipid order in the presence and absence of recombinant EBOV VP40. The ratio of probe:lipids was 1:200 molar ratio, and the excitation wavelength was 430 nm. E) Representative confocal images of fluorescent GUVs (green) with different lipid orders and compositions containing 20% PS and 5% PI(4,5)P_2_ molar ratio, were incubated with 1.25 µM of EBOV VP40:VP40-AlexaFluor 594 (9:1 molar ratio). The white arrowheads indicate membrane tethering induced by protein enrichment and white arrows indicate regions on the GUV membrane that were able to enrich uniformly with VP40 without tethering. F) Enrichment factor of VP40 on the GUV membranes according to the different lipid ordering states determined by the ratio of total fluorescence intensity of the protein over the total analyzed area of the GUV membrane. The values are reported as mean ± SEM of three independent replicates. G) Representative SPR binding response curves of VP40 to LUVs containing different species of PS (22% molar ratio) according to their fatty acid tail saturations; DOPS: dioleoyl-PS; POPS: palmitoyl oleoyl-PS; DPPS: dipalmitoyl-PS. The apparent affinity of VP40 for the different PS-containing membranes was determined by fitting the maximum response obtained from three sets of SPR binding curves vs different concentrations of VP40 to a Langmuir binding isotherm (6, 8).

To determine if VP40 can alter the membrane fluidity of a simplified *in vitro* lipid vesicle system, we used a push-pull pyrene (PA) probe (28) for assessing membrane fluidity/rigidity. The PA probe is similar to the Laurdan dye with strong brightness and much higher photostability in the non-polar membrane environments (28). VP40 membrane binding increased exclusively the rigidity of the S_o_ membranes to yield a similar spectrum as the L_o_ membranes with or without VP40. In contrast, L_d_ membranes did not display a detectable shift in spectrum. Two types of PI(4,5)P_2_ were utilized for these measurements (18:1 and brain PIP_2_) to assess if fluidity changes of the membrane were due to VP40 binding to PI(4,5)P_2_ and potential clustering of this lipid upon VP40 matrix formation. Brain PIP_2_ has a high amount of polyunsaturated (18:0, 20:4) species (57). The PA probe fluidity assay showed the increase of rigidity observed in S_o_ membranes was independent of the PIP_2_ species assessed; indicating rigidity increases *in vitro* were independent of the PIP_2_ acyl chain (Fig. 3D).

To further investigate if the fatty acid environment within the membrane leaflets is involved in the association and enrichment of VP40 on membranes, we performed confocal imaging of giant unilamellar vesicles (GUVs) with different acyl chain compositions. First, we conjugated Alexa Fluor-594™ C5 Maleimide on the cysteine residues on the C-terminal of VP40 (C311/C314) (11). A previous functional study on the effect of alanine substitution of these residues showed a slight increase in binding of VP40 to PS containing vesicles, however, mutations of C311 and/or C314 had no significant effect on VP40 PM localization (58). The protein was added at a ratio of 9:1 (VP40:VP40-Alexa Fluor-594) to the vesicle solution and incubated for 30 min at 37°C prior to imaging. The fluorescence signal enriched on membranes to different extents depending on the fatty acid tail composition and rigidity states of the membranes (Fig. 3E). A more rigid and ordered hydrophobic environment within the GUVs led to an increase of VP40 enrichment (Fig. 3E-F) confirming our previous observations from the membrane pelleting assay (Fig. 3B-C). The fluorescence was more evenly distributed on the membrane with increased membrane rigidity (Fig. 3E, arrows). Furthermore, we observed tethering of GUV membranes with a preferential accumulation of VP40 at GUV-GUV contact sites (Fig. 3E, arrowheads). In the absence of VP40 on the GUV membranes, we did not observe GUV-GUV membrane adhesion. In all cases, membrane deformation was observed in clustered GUVs and sometimes membrane bending and vesiculation were observed, however, no membrane scission, hemifusion, or full fusion were detected during the imaging time frames studied (Fig. S3A-B).

Since PIP_2_ fatty acid moieties don’t seem to be involved in VP40-induced membrane fluidity changes, we compared how the fatty acid tails of different PS species influence the binding affinity of VP40 using a surface plasmon resonance (SPR) assay (6, 8). In the previously studied membranes, the lipid compositions between L_d_, L_o_, and S_o_ membranes were different for the purpose of the assays. For this experiment, we conserved the same lipid backbone (POPC) which was held constant at a 78% molar ratio and PS with different acyl chain moieties was changed at a 22% molar ratio. The equilibrium dissociation constant (*K*_d_) from this analysis indicated a thirty- and eighteen-times greater apparent binding affinity to DPPS (10 nM) compared to DOPS (311 nM) and POPS (186 nM), respectively (Fig. 3G). Taken together, the data described here show that the acyl chain composition of the PS species endow increased ability of VP40 to bind membranes, where VP40 shows binding selectivity for both lipid head groups and lipid acyl chain composition.

### PM fluidity impairment results in VLP retraction, budding inhibition and reduced viral infectivity

To evaluate if the PM fluidity changes can be a target to reduce EBOV budding and infectivity, we first performed transmission electron microscopy (TEM) on cells expressing VP40 and treated with dibucaine, propranolol or TFP to investigate the structure of VLPs at the surface of cells. Fig. 4A shows that 30 minutes post-treatment, a significant reduction of the density of VLPs at the surface of cells was observed with seemingly shorter lengths compared to VLPs observed on the surface of non-treated cells (Fig. S4A).

**Figure 4.**
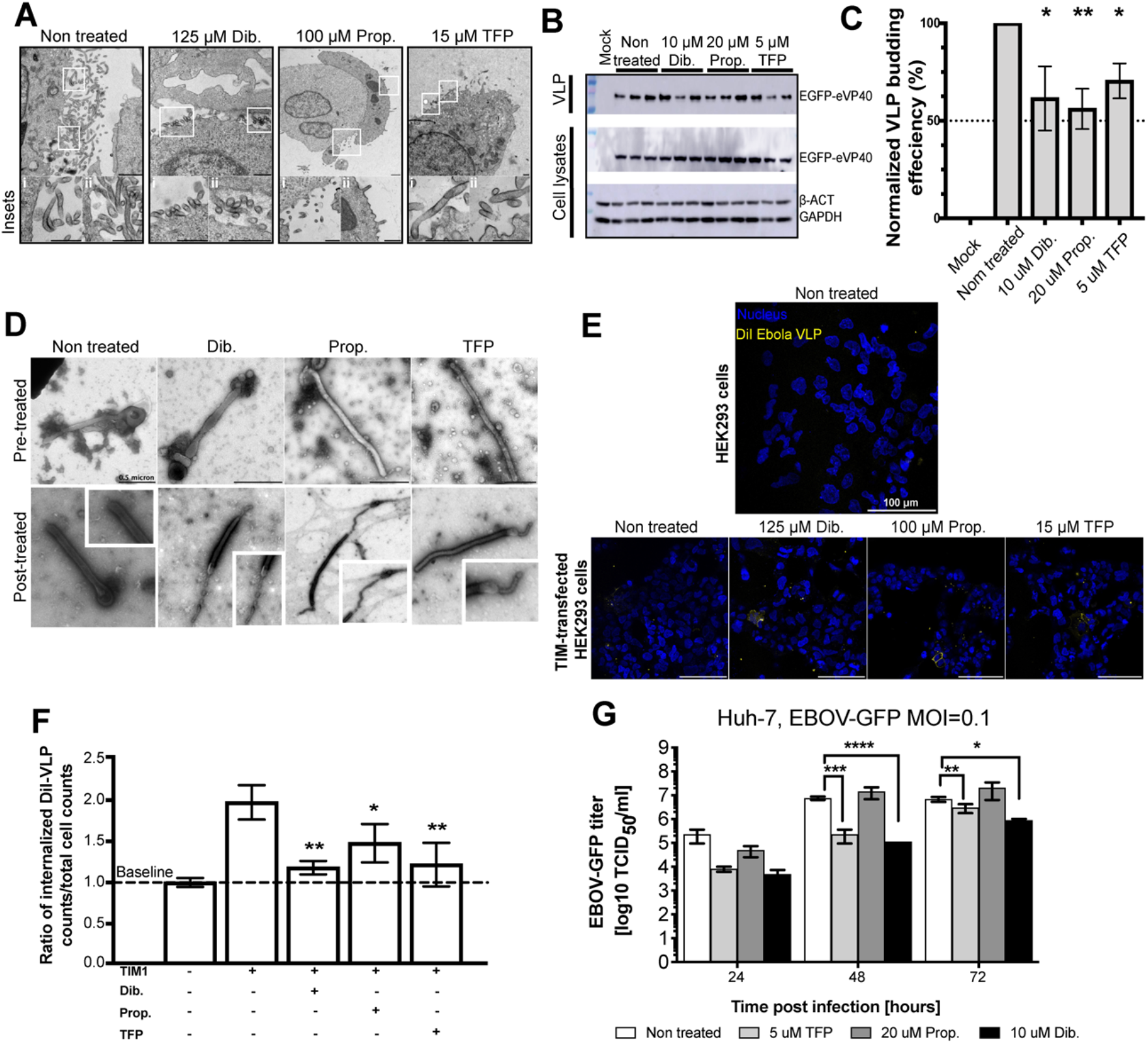
Membrane fluidity impairment of EBOV infectivity by disrupting viral matrix assembly and the VLP integrity. A) Representative TEM micrographs of HEK293 cells expressing VP40 treated with corresponding drugs to fluidize the PM for 30 min prior to processing. The zoomed insets show reduced filamentous structures at the surface of treated cells compared to non-treated control cells. B) Immunoblot of VLP budding assays performed 72 hours post-transfection. HEK293 cells expressing VP40 were treated with the different FDA approved drugs 16 hours post-transfection and the conditions were maintained until VLP collection. C) The relative budding efficiency of VP40 VLPs under the indicated conditions in B) from three independent experiments reported as mean ± SEM. One-way ANOVA with multiple comparisons were performed, \**P*,0.003, \*\**P*<0.003. D) Representative TEM micrographs of negatively stained VLPs pre- and post-treated with dibucaine, propranolol or TFP post-collection. The zoomed in insets in the treated VLP images highlight the morphological defects induced by the treatments. E) Representative Z-stack reconstituted confocal images of a fluorescence based DiI TIM-1 dependent VLP entry assay. HEK293 cells expressing TIM-1 or not (target cells) were incubated with non-treated, or drug treated VLPs pre-stained with DiI (yellow). The DNA of cells is stained (blue) and the number of VLPs per cell (total number of yellow puncta/total number of nuclei) was determined as a ratio of internalized VLP as shown in (F). The values were normalized where the absence of TIM-1 in target cells incubated with non-treated VLPs was considered as a baseline for the entry assay. The values are reported as mean ± SD. One-way ANOVA with multiple comparisons were performed, \**P*,0.003, \*\**P*<0.003. To evaluate the effect of drugs on authentic EBOV infectivity, Huh-7 cells were infected with EBOV-GFP at a multiplicity of infection (MOI) of 0.1. Viral titers were determined over time (G). Experiments were carried out in three independent replicates and two-way ANOVA analysis with Tukey multiple comparisons performed (**** *P*<0.0001; *** *P*<0.001).

We next evaluated the ability of VP40 expressing cells to release VLPs and the ability of these VLPs to enter cells. A VLP budding assay was first performed on cells treated 5 hours post-transfection with VP40 and the treatment with drug was maintained for 72 hours. For this assay, the drug concentrations were reduced to limit the cytotoxic effects of dibucaine (IC_50_ = 56 µM), propranolol (IC_50_ = 57.7 µM) and TFP (IC_50_ = 36.7 µM) to 10 µM, 20 µM and 5 µM, respectively. The quantitative analysis indicated a 30 to 45% reduction in VLP production upon drug treatment (Fig. 4B-C). To evaluate the off-target impact of these FDA-approved drug treatments on protein synthesis that can lead to reduced VLP production, we estimated the total protein content per viable cell during the treatments (every 24 hours). The analysis showed no significant difference among the experimental conditions (Fig. S4B) and the average protein mass/cell for non-treated cells (0.24 ± 0.04, 0.17 ± 0.04 and 0.18 ± 0.02 ng/cell for 24, 48 and 72 hours, respectively) were comparable to previously reported data (0.18 ± 0.01 ng/cell) (59) suggesting that protein synthesis in the treated cells is not appreciably affected by the drugs and reduction of VLP release is due to the inability of VP40 to properly assemble at the PM.

To further support our observations, we performed negative stain and TEM on fixed VLPs collected from cells treated with the FDA-approved drugs. First, the average lengths and diameters of VLPs collected from non-treated cells, dibucaine, propranolol and TFP treated cells were very similar, 70 nm ± 15 and 1.7 nm ± 1.8 (Fig. S4C-D), respectively. Next, we analyzed if the drugs affected the VLP structural integrity that may have been sensitive to our collection method. To test this hypothesis, we collected VLPs 72 hours post-transfection, then treated the VLPs for 30 min with the different compounds. Here we observed serious damage of the VLP lipid envelopes resulting in naked segments of the VLPs. This indicated that the FDA-approved drugs can also affect the fluidity and integrity of the lipid membrane of the VLP (Fig. 4D, bottom panel).

The abolished structures of the VLPs in addition to the reduced amount of VLPs from drug treated cells lead us to investigate the ability of VLPs to enter cells over-expressing T-cell immunoglobulin and mucin domain 1 (TIM-1) receptor. To test the VLP entry ability, EBOV glycoprotein (GP) was co-expressed with VP40 to produce entry competent VLPs. VLPs were stained with the lipophilic DiI stain post collection (10), then treated with corresponding drugs for 30 min prior to cell inoculation. Confocal imaging on cells post-entry showed that over-expression of TIM-1 lead to a two-fold increase in GP-containing VLP entry (Fig. 4E-F, (60, 61)). In contrast, VLPs had 45-50% reduced entry efficiency when treated with dibucaine and TFP, while propranolol reduced the amount of VLP internalized by ∼25% (Fig. 4E-F). These data confirm that the structural damage induced upon drug treatment reduced VLP ability to enter and infect cells as well the potential altered pools of PS for TIM-1 interactions. Further, the effect of the fluidity increases on the GP distribution and/or re-organization on the viral envelope is not known and could contribute to alterations in the interactions of GP with host proteins.

Our investigations described above were conducted outside of the context of authentic virus. Thus, we investigated if impairment of lipid membrane fluidity by the FDA-approved drugs would also affect infectivity of authentic EBOV. We performed virus growth kinetics using EBOV-GFP (62) in cells treated with the drugs post-infection. Supernatant samples were collected every 24 hours for 3 days and the virus titers were determined (Fig. 4G). The most significant reduction of viral titers was observed at 48 hours post-infection with treatment for both dibucaine and TFP (Fig. 4G, Fig. S5). This effect was also observed at 72 hours but with less significant differences. In contrast, propranolol exhibited an insignificant reduction of titers at 24 hours but no effect after 48- and 72-hours post-infection. Taken together, these data show that the membrane fluidity is an important factor to stabilize the VP40 matrix assembly, a required step for virus budding, integrity and infectivity providing new perspectives for therapeutic development to prevent viral spread.

## Discussion

During viral particle assembly, EBOV VP40 associates with the PM inner leaflet (63) where the viral GP trimers are already integrated. The co-expression of the GP with VP40 enhances VLP egress 5-8 fold (64-67). It is well-established that PM phospholipid composition of the anionic lipids PS and PI(4,5)P_2_ are critical components to trigger peripheral VP40 membrane association. PS is proposed to be involved in the initiation of small oligomer formation (6, 9, 68), and PI(4,5)P_2_ involved in clustering and stabilizing large oligomers to form the viral matrix (7). Also, VP40 large oligomers were observed to associate with lipid rafts (39). However, cholesterol levels showed no impact on the ability of EBOV VP40 to properly associate to PS-containing membranes (69). For Marburg virus (MARV), VP40 displays a more relaxed specificity to PS and PI(4,5)P_2_ where it can interact with different anionic phospholipids in a nonspecific electrostatic fashion but PS and PIP_2_ are required for the protein to undergo proper oligomerization through NTD-NTD and CTD-CTD, respectively (70, 71).

In contrast to EBOV VP40, MARV VP40’s binding to lipids is enhanced in the presence of cholesterol (70). The cholesterol in phospholipid membranes increases the packing density of the lipid headgroups and acyl chains leading to an increased acyl chain order (72). The difference in behavior of EBOV and MARV VP40 towards lipid membranes make for excellent model systems *in vitro* and in cells to probe and understand the biophysical mechanisms involved in filovirus matrix assembly. Besides the PM anionic charge (PS and PI(4,5)P_2_ specifically), little is known of other factors of the PM that make it more attractive to EBOV VP40 and that are required for an efficient and stable matrix formation. In this study, we aimed to deepen our fundamental understanding of EBOV VP40 membrane binding and matrix assembly and to potentially advance novel therapeutic strategies.

Cell membrane fluidity is critical to maintain membrane function as well as protein and lipid dynamics through the bilayer. Herein we report the ability of EBOV VP40 to increase PM fluidity. Active lipid flip-flop and scrambling and lipid lateral diffusion can lead to membrane fluidity increases. EBOV VP40 binding and assembly results in PS exposure to the outer leaflet of the PM and viral envelope (6) by activating the scramblase XK-related protein 8 (Xkr8) (10, 11). This externalization of PS is exploited by EBOV as an apoptotic mimicry strategy to facilitate the viral entry (73-75). Furthermore, EBOV VP40 clusters fluorescent labelled PS at the budding site of the PM (12). The ability of VP40 to bend membranes can lead to the recruitment of host factors including conical shaped lipids to promote membrane curvature and recruitment or activity of membrane scission machinery. Conical lipids include polyunsaturated fatty acid-containing phospholipids and glycerolipids (76). The accumulation of these lipids at the PM can also result in the increase of membrane fluidity, favorable to positive membrane curvature.

The lipidomic analysis of serum from patients that survived EBOV disease (EVD) compared to fatalities and a healthy control group, highlighted an overall increase of PS abundance in fatalities (77). In addition, EVD PS signatures suggested an increase of PS(16:0;18:1) and (16:0;22:6). Extracellular vesicles are enriched in PS lipids, especially PS(18:0_18:1), PS(18:0_20:4) and PS(18:0_22:6) (77-79). The PS ratio of PS(18:0_20:4) divided by the sum of PS(18:0_18:1), PS(18:0_20:4) and PS(16:0;22:6) trends to be lower in EBOV fatalities compared to the healthy group (77). These severe changes in the PS species are consistent with the regulation of PS metabolism, trafficking and translocation previously reported upon EBOV infection or VP40 over-expression (6, 10, 73).

We reported the importance of an ordered hydrophobic environment for stable binding and oligomerization of VP40. The three FDA-approved drugs tested in this study accumulate within the lipid leaflets. We also showed that all three drugs have a reversible effect on the plasma membrane fluidity and VP40 matrix assembly. We can speculate that at 72 hours post-infection and drug treatment start, the effect of these drugs starts to fade. Thus, continuous treatment or drug replenishment every 24 or 48 hours may result in a higher reduction of viral titers. Dibucaine accumulates within the membrane leaflets and makes favorable electrostatic interactions with lipid glycerol groups and hydrophobic contacts with acyl chains (80). Furthermore, PS is involved in dibucaine’s quinoline ring interaction with Phe residues of the channel target and can be externalized in platelets as mitochondrial apoptotic-like response to dibucaine treatment (even in calcium-free conditions) (80, 81). In a simplified lipid vesicle system, dibucaine reduced the miscibility temperature of lipids and the line tension between the different phases (L_o_ and L_d_) resulting in disordered lipid packing and increased fluidity of L_o_ phase-membranes (37, 80). The normal binding of LactC2 to the PM upon high concentration dibucaine treatment supports that the observed defect in VP40 binding to the PM is due to an increase in membrane fluidity. (33, 82). The β-blocker, propranolol, increases the membrane fluidity by lowering the melting temperature of lipids and decreasing the phase transition within the membrane (33). Propranolol aligns perpendicular to the membrane bilayer near the membrane surface (82, 83). It exhibits more contacts with atoms in the headgroup and glycerol interfaces which perturbs the proper alignment of phospholipids resulting in fluidizing the membrane (33, 82). This suggests that VP40 binding stability may also be sensitive to the distance between the lipid headgroups.

Lipid-raft domains are the assembly and budding platforms of several enveloped viruses including Influenza A, HIV-1, and paramyxoviruses. The HA of Influenza A virus and GP of EBOV and MARV as well as measles membrane associated proteins (F, M and H) have been shown to be enriched in DRM-raft domains (85-88). In addition, the oligomerization of peripheral proteins and assembly of these viruses has been reported to occur in DRM of infected or transfected cells (39, 88-94). In the case of filoviruses, GP is mostly incorporated in DRM and both VLPs and authentic virus contain raft resident proteins such as GM1 as observed by confocal imaging and immunoprecipitation of filoviruses particles (88). Liquid chromatography-tandem mass spectrometry analysis of EBOV and MARV particles were consistent with previous results (88), where half of host proteins identified by LC-MS/MS are described as raft resident proteins (95). In alignment with these observations, DRM are mostly enriched of multimerized VP40 (39) as we also observed in this study. Altogether, our observations and prior literature highly support the importance of DRM in EBOV assembly. The current study demonstrated VP40’s attraction towards DRM is most likely due to the highly ordered state of these membranes independent of cholesterol enrichment.

EBOV infection results in hemorrhagic disease with high fever. The increase of body temperature may affect the fluidity of cellular membranes, however, from this current study we know that the viral matrix assembly is sensitive to an increase in membrane fluidity. Therefore, we propose a model (Fig. 5) in which the incorporation of GP within DRM may improve and/or stabilize the order within the leaflets of these membrane domains. Since GP co-expression with VP40 enhances VLP budding (64, 84), this may aid in viral matrix assembly and release during the infection to ensure efficient viral production.

**Figure 5.**
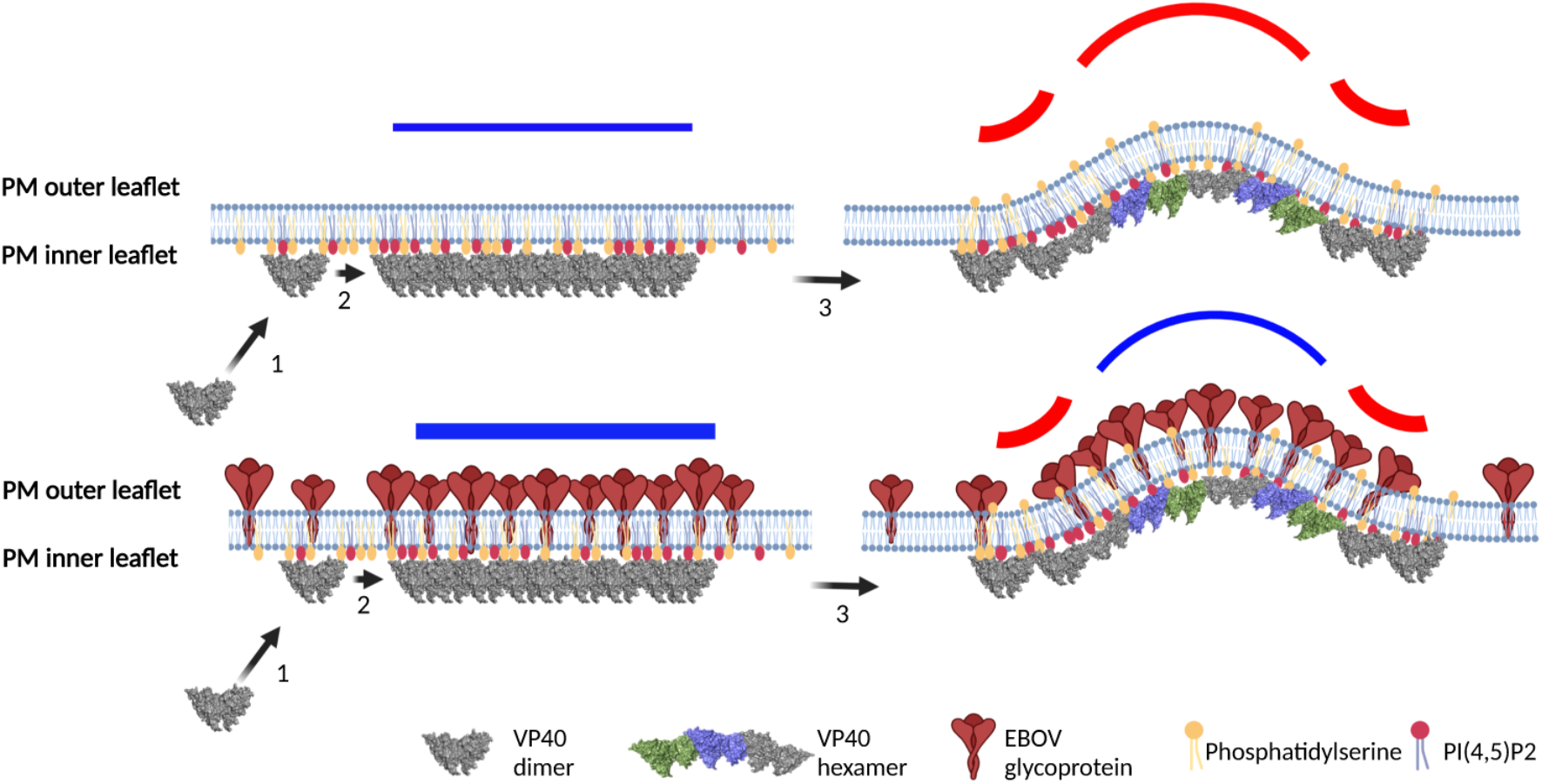
Model depicting the proposed effect of membrane rigidity on EBOV VP40 matrix assembly. In absence of glycoprotein (GP) (upper panel), EBOV VP40 (1) traffics to the PM independent of membrane rigidity, (2) binds to the most abundant anionic lipid in the PM inner leaflet, PS (6, 8, 58), to initiate the matrix assembly (dimer to hexamer). PM binding of VP40 is supported by the interaction of VP40-PI(4,5)P_2_ (7), a key lipid for the extensive oligomerization of the viral protein and matrix formation (63). This extensive assembly leads to lipid clustering (12), PS exposure on the outer leaflet (10, 11) and membrane curvature to initiate viral budding. These VP40-induced events result in an increase in PM fluidity at the assembly and budding sites. In the presence of GP (lower panel), we propose that GP increases the order and organization of the hydrophobic environment within the PM leaflet. This likely promotes VP40 recruitment at the PM, stabilizes the viral matrix formation and may aid in maintaining the PM rigidity during viral assembly. Schematic created with BioRender.

The ability of VP40 to tether membranes and induce the clustering of GUVs was consistently observed. We also noticed that the bound VP40 within the tethering interface made the membranes flatter and in some cases membrane bending was also observed without any detectable membrane fusion. We speculate that the forces on both sides of the tethering interface may be equal suggesting a similar tight binding to the lipids may be occurring on each membrane and the tethering a direct consequence of the VP40 oligomerization ability (Fig. S3C). Currently, two models of VP40 matrix assembly have been proposed (63, 96). One model promotes VP40 hexamers formed from rearranged dimers as a building block of the matrix (96), while a more recent model proposes a matrix lattice formed from a patchwork of VP40 dimers assembled via CTD-CTD interactions across the membrane (63). In the model in Fig S3C, we assume that on each side of the tethering interface, VP40 interacts with the membranes through its C-terminal domain. In cells, EBOV VP40 is typically observed diffuse in the cytosol or accumulated at the PM inner leaflet. VP40 protein organization and rearrangement at cellular membranes still needs to be further determined. For this reason, we believe the capacity of VP40 to tether membranes may occur close to or at the PM interface as a result of VP40 trafficking on carrier vesicles.

The ER is the key site of phospholipid biosynthesis and close contacts between organelle membranes and the ER facilitates transfer of lipids between organelles (97-102). Due to the large demand on PS during EBOV assembly and release at the PM, we hypothesize that the VP40 tethering capacity may help to promote the contact between ER and the PM (98). However, no evidence is available in the literature about such a phenotype occurring during EBOV infection or VP40 transfected cells. EBOV VP40 trafficking to the plasma uses the COPII transport system (103). This system works at an initial step of the secretory pathway in ER-Golgi transport and has not been described to be involved in cargo trafficking to the PM. However, VP40 relies on this system to reach the PM through its interaction with Sec24C (103, 104), an essential component of COPII vesicles. The mechanism used by VP40 to hijack this system is still unclear. It is possible that the ability of VP40 to tether membranes is involved in promoting its proper trafficking to the PM and its interaction with cellular factors such as Tsg101 or Nedd4, key factors for the recruitment of the ESCRT machinery, required to initiate viral budding from the PM (2, 105). EBOV GP accumulates in the ER of infected or transfected cells and is transported to the PM (106-108). The co-expression of VP40 and GP did not change GP cellular distribution (106). Another hypothesis is that VP40-mediated membrane tethering may facilitate the translocation of GP to the PM. Further investigations of membrane tethering mediated by VP40 within the cell will greatly improve our understanding of the VP40 function in cells.

In summary, our study highlights new properties of the PM including an ordered hydrophobic environment required for EBOV VP40 to bind stably, oligomerize, assemble new viral particles and conserve the infectious morphology of released particles. These findings greatly enhance our understanding of the molecular and biophysical mechanisms of EBOV assembly and suggest new directions for further investigations and for development of novel therapeutic strategies against EVD.

## Materials and Methods

### Pharmaceutical Treatments

Stock solutions of dibucaine-HCl (500 mM in ddH_2_O), propranolol-HCl (125 mM in ddH_2_O) and trifluoperazine-2HCl (50 mM in ddH_2_O) were prepared. For confocal imaging experiments and N&B experiments, cells were treated for 5 minutes with 1 mM dibucaine, 1 mM propranolol, or 0.1 mM trifluoperazine in calcium free media (150 mM NaCl, 5 mM KCl, 1 mM MgCl_2_, 20 mM HEPES, 100 μM EGTA, pH 7.4). For Laurdan fluidity assays, membrane fractionation assays, TEM, and SEM experiments and VLP treatments, samples were treated for 30 minutes with 125 µM dibucaine, 100 µM propranolol, or 15 µM trifluoperazine in calcium free media. For VLP budding assays and virus titer experiments, cells were treated with 10 µM dibucaine, 20 µM propranolol, or 5 µM trifluoperazine in calcium free media as described in the respective method sections. Methyl-β-cyclodextrin (MβCD, Sigma-Aldrich, C4555) was prepared in 1X PBS, pH 7.4 and cells were treated with 10 mM methyl-β-cyclodextrin for 30 minutes prior to imaging. 12:0 dihydrosphingomyelin (DH-SM (d18:0/12:0), Avanti Polar Lipids, 860683) was prepared in ethanol and used to treated HEK293 cells at a final concentration of 2.5 μM for 3 hours at 37°C (adapted from (109)). Laurdan treatments were performed as described previously (71). Primary antibodies used for Western blotting were rabbit anti-EBOV antibody (IBT Bioservices, 0301-010), mouse anti-EBOV GP antibody (IBT Bioservices, 0365-001), HRP-conjugated mouse anti-GAPDH antibody (Fisher Scientific, CB1001), HRP-conjugated mouse anti-β-actin antibody (Santa Cruz Biotechnologies, sc-47778), mouse anti-Caveolin 1 antibody (Cell Signaling Technology, 3238S) and HRP-conjugated rabbit anti-GFP antibody (Abcam, ab190584). Secondary antibodies used for western blotting included HRP-conjugated goat anti-Rabbit and sheep anti-Mouse antibodies (Abcam, ab205718 and ab6808, respectively).

### Virus-like particle entry assay

VLPs were produced from HEK293 cells (5 × 10^6^ cells) expressing VP40 and GP were purified and stained with DiI according to (10) and as described in supplementary information. After staining, 1x Halt protease inhibitor cocktail was added to the VLP collected in phenol-free MEM with 2% HD-FBS and 4% BSA. For each condition, 120 µl of VLP fractions were treated, for 30 min on ice and in the dark, with 1 µl of FDA approved drugs for a final concentration of 125 µM dibucaine, 100 µM propranolol, 15 µM trifluoperazine diluted in the same media as above. To assess VLP entry, HEK293 cells were transfected with TIM-1 for 24 hours prior to incubation with DiI-labeled VLPs and briefly rinsed with phenol-free MEM with 2% HD-FBS and 4% BSA. DiI-VLPs were added to TIM-1 expressing HEK293 cells, spinoculated for 45 min at 4°C, and allowed to incubate for 1 hour at 37°C. Plates were then rinsed with PBS and nuclei were stained with Hoechst 3342 for 15 min at 37 ºC prior to fixation with 4% paraformaldehyde (PFA) and confocal imaging.

### Confocal imaging

Live cell imaging experiments were performed 24 hours post-transfection and 30 min post-treatments. Confocal imaging was performed to quantify plasma membrane localization of VP40 or control proteins using either a Zeiss LSM 710 or a Nikon Eclipse Ti Confocal inverted microscopes using a Plan Apochromat 63× 1.4 numerical aperture oil objective. A 488 nm argon laser was used to excite GFP/EGFP and a 561 nm diode laser was used to excite RFP and mCherry.

To assess Laurdan fluidity assay at the plasma membrane of cells expressing EBOV VP40, we followed the protocol as described previously (71, 110). In brief, cells were treated with 10 μM Laurdan for 30 min at 37°C. A minimum of ten cells per replicate were imaged with a multi-photon confocal microscope. The dye was excited at 800 nm, and emission intensities were recorded simultaneously in the 400–460 nm and 470–530 nm ranges with photon multiplier tubes (PMTs) for ordered (PMT1) and disordered (PMT2) membranes, respectively. Calibration images were acquired with 100 μM Laurdan in culture media to calculate the measured generalized polarization factor. Image processing was done using ImageJ, and generalized polarization index distribution was determined using the Laurdan_GP macro (110).

### Membrane fractionations for DRM and DSM isolation

HEK293 cells (2 × 10^7^) expressing EGFP-VP40 for 72 hours were treated with 125 µM dibucaine, 100 µM propranolol, 15 µM trifluoperazine for 30 minutes at 37 ºC. Cell lysis and DRM/lipid raft and DSM/non raft membranes were generated as described previously (39, 40). 25 fractions of 200 µl, in addition to the pellet fraction corresponding to the cytoskeleton were collected and separated on 10% acrylamide gels using SDS-PAGE without boiling samples prior to loading. The oligomerization and subcellular distribution of VP40 were then determined by Western blotting.

### EBOV growth kinetics

All work involving EBOV was performed in the maximum containment laboratory (MCL) at the Rocky Mountain Laboratories (RML), Division of Intramural Research, National Institute of Allergy and Infectious Diseases, National Institutes of Health. Huh-7 cells (mycoplasma negative) were seeded in 24 well plates the day before infection. Cells were infected (5 × 10^5^ cells/well) with EBOV-GFP at a MOI=0.1 (5 × 10^4^ ffu/well in 0.5 ml) for 1 hour. The inoculum was removed, cells were washed twice with plain DMEM, and 1.5 ml Ca^2+^-free media (HBSS) containing the drugs were added. Cells were monitored for GFP expression (111) and samples were collected (100 µl per time point) for virus titration at 0, 24, 48, and 72 hours. The median tissue culture infectious dose (TCID_50_) was determined from each sample as previously described using the Reed and Muench method (112).

### GUV formation and imaging

GUVs were prepared by a gentle hydration method (113–115). Briefly, 1 mM of lipid control mixtures were made and contained: *DOPLs* (1,2-dioleoyl-sn-glycero-3-phosphocholine (DOPC), 1,2-dioleoyl-3-trimethylammonium-propane (DOTAP) and TopFluor® fluorescent lipid (89.8:10:0.2% molar ratio)); *POPLs* (1-palmitoyl-2-oleoyl-glycero-3-PC (POPC), 1-palmitoyl-2-oleoyl-glycero-3-phosphoethanolamine (POPE) and TopFluor® fluorescent lipid (89.8:10:0.2% molar ratio)); *DPPLs* (1,2-dipalmitoyl-sn-glycero-3-PC (DPPC), cholesterol (Chol) and TopFluor® fluorescent lipid (49.9:49.9:0.2% molar ratio)). Different PS species and brain phosphatidylinositol 4,5-bisphosphate PI(4,5)P_2_ were added in lipid mixtures at %molar ratios of 40% and 5%, respectively to DOPLs (DOPS), POPLs (POPS) and DPPLs (DPPS) and the ratios of (i) phosphatidylcholine (PC) were adjusted accordingly in DOPLs and POPLs, while in (ii) DPPLs, the ratios of both PC and Chol were adjusted to maintain a total %molar ratio of 100%. The lipid mixtures were prepared in 5 mL round-bottom glass flasks and the chloroform was removed with rotary movements under a continuous stream of N_2_ and even heat at 55°C (DPPLs mixtures) until complete evaporation of the solvent. The lipid films were then hydrated overnight at 37°C (DOPLs and POPLs) and 55°C (DPPLs) in an appropriate volume of GUV hydration buffer (150 mM NaCl, 10 mM HEPES, 0.5 M sucrose, pH 7.4).

For imaging, the freshly hydrated GUVs were diluted 10 times in GUV dilution buffer (150 mM NaCl, 10 mM HEPES, 0.5 M glucose, pH 7.4) and placed on a 6-mm diameter chamber made from a silicon sheet using a core sampling tool (EMS, 69039-60). The silicon chamber was mounted on a 1.5-mm clean cover glass (EMS, 72200-31) pre-coated with 1 mg/mL BSA. The set up was then assembled in an Attofluor chamber (Invitrogen, A7816) and VP40-Alexa594 was added and incubated with lipids for 30 min at 37°C. GUV imaging was performed at 37°C for no longer than 30 min on a Nikon Eclipse Ti Confocal inverted microscope (Nikon Instruments, Japan), using either a Plan Apochromat 60x 1.4 numerical aperture oil objective or a 100x 1.45 numerical aperture oil objective, respectively. A 488 nm argon laser was used to excite TopFluor® lipids and a 561 nm argon laser was used to excite eVP40-Alexa594. Protein enrichment at the GUV membranes was estimated using Image J (116), as follows:

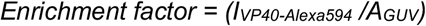

Where I corresponds to the intensity of VP40-Alexa594 at the membrane of the GUVs after background subtraction, and A correspond to the area covering the GUV membranes.

## Acknowledgments

S.A. and R.V.S. thank Nathan J. Dissinger for excellent technical support. The authors acknowledge the use of the facilities of the Bindley Bioscience Center, a core facility of the NIH-funded Indiana Clinical and Translational Sciences institute, the use of the Purdue Life Science Electron Microscopy facility, and the use of the Integrated Research Facility at Rocky Mountain Laboratories, Division of Intramural Research, NIAID, NIH. These studies were supported by the NIH AI0817 to R.V.S., NIH T32 GM075762 to M.L.H. and the Purdue Pharmacy Live Cell Imaging Facility (NIH OD027034 to R.V.S.). A.M. is supported by the Division of Intramural Research, NIAID, NIH.

## Supporting information

### Material and Methods

#### Cells and Plasmids

HEK293, Huh-7 and COS-7 cells were cultured in Dulbecco’s modified Eagle’s medium (DMEM) supplemented with 10% heat-inactivated fetal bovine serum (FBS) and 1% penicillin/streptomycin. Cell transfections were performed using Lipofectamine 2000 according to the manufacturer’s instructions. Plasmids coding for EGFP and EGFP-VP40 were prepared as described previously (1, 2). The GFP-LactC2 plasmid was a kind gift from Sergio Grinstein (University of Toronto). GFP-PLCδ-PH was a kind gift from Tamas Balla (NIH). pcDNA3.1-eGP (NR-19814) was obtained through BEI Resources (NIAID, NIH). pCAGGS-TIM-1 was provided by Heinz Feldmann (3). The plasmid encoding the full-length of EBOV VP40 pET46-VP40-6xHis (pET46 Ek/LIC) DNA was a kind gift from Erica Ollmann Saphire (La Jolla Institute for Immunology); pCEboZVP40 from Dr. Kawaoka (4), and KRphi-mRFP a gift from Sergio Grinstein (Addgene plasmid # 17276).

#### Virus-like particle (VLP) production and budding assay

HEK293 cells (2.5 × 10^6^) were transfected with EGFP-VP40 plasmid in a 6-well plate. Drug treatments were started 16 hours post-transfection using fresh complemented media. The treatments were maintained for 72 hours until VLP collection. To purify the VLPs, the clarified cell supernatants from treated cells were pelleted over a 20% (wt/vol) sucrose cushion as previously described (5). VLP samples were stored at - 80°C until further analyses. The budding efficiency was determined by Western blotting (2, 6). All quantitative analysis derived from western blotting was performed using densitometry analysis in ImageJ (7). The following equation was applied:

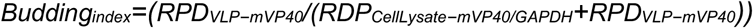

RPD is the relative pixel density. The budding index of each mutant was normalized to the non-treated budding index.

#### Total internal refection fluorescence (TIRF) imaging

HEK293 cells were seeded and transfected with EGFP-VP40 as described above on 40 mm extra clean coverslips. 24 hours post-transfection, the 40 mm coverslips were assembled into a RC-31 imaging chamber (Warner instruments, Hamden, CT) and attached to a six-channel perfusion mini-valve system (VC-6M). Cells were then washed for 10 min with calcium free media for equilibration, then treated with different FDA-approved drug concentrations alternated with 8 to 10 min calcium free media washes. The cells were imaged to assess the effect of the drugs on VP40 binding to the plasma membrane using a Nikon A1 Confocal and Perfect Focus Ti-E Inverted Microscope equipped with an Apo TIRF 60x Oil DIC N2 (NA 1.49) objective lens.

#### Transmission electron microscopy (TEM)

HEK293 cells were transfected with EGFP-VP40 and treated as described above. 24 hours post-transfection, cells were washed, fixed with, and stored in cold primary fixative (2.5% glutaraldehyde in 0.1 M sodium cacodylate buffer), rinsed, and embedded in agarose. Small pieces of cell pellet were post-fixed in buffered 1% osmium tetroxide containing 0.8% potassium ferricyanide and en bloc stained in 1% uranyl acetate. They were then dehydrated with a graded series of ethanol, transferred into acetonitrile, and embedded in EMbed-812 resin. Thin sections were cut on a Reichert-Jung Ultracut E ultramicrotome and stained with 4% uranyl acetate and lead citrate. Images were acquired on a FEI Tecnai T12 electron microscope equipped with a tungsten source and operating at 80 kV.

Purified VLPs were fixed with 2.5% glutaraldehyde in 0.1 M cacodylate buffer. Purified VLPs were applied onto glow discharged carbon formvar grids and negatively stained using 2% phosphotungstic acid solution (PTA). Samples were imaged with the same electron microscope than above. VLP length and diameter measurements were quantified using ImageJ software. For diameter analysis, multiple different diameters were measured across random areas on each VLP, and the mean diameter was reported.

#### Scanning electron microscopy (SEM)

HEK293 cells were transfected with EGFP-VP40 and treated as described above. Cells were scraped and collected through low-speed centrifugation at 48 hours post-transfection, and stored in primary fixative (2% glutaraldehyde, 2% paraformaldehyde in 0.1 M cacodylate buffer, pH 7.35) at 4°C until processing. During processing, samples were fixed to coverslips and post-stained with 1% osmium tetroxide in 0.1 M cacodylate buffer. Samples were extensively rinsed with water and dehydrated with a graded series of ethanol followed by drying in a Tousimis 931 Supercritical Autosamdri® device. Prior to imaging, samples were coated with 3 nm Iridium. A Field Emission Scanning Electron Microscope Magellan 400 (FEI) (Hillsboro, OR) was used to collect images, with assistance from Tatyana Orlova at the Notre Dame Integrated Imaging Facility.

#### Number and Brightness (N&B) assay

VP40 oligomer size was measured before and after FDA approved drug treatments. This method was carried out as previously described (8–10). Olympus FluoView FV1000 microscope with a 40X 1.3 numerical aperture oil objective was used for data collection: 100 images taken with raster scanning with a 12.5 μs pixel dwell time, zoom set to 16.4, 256 × 256 frame size, photon count mode on. Image analysis was performed in SimFCS software. The brightness of EGFP transfected into COS-7 cells was determined as previously reported (2, 11). The ratio of pixels shown as a monomer-hexamer, hexamer-12mer, and 12mer + were calculated for each before and after treatments. The before treatment values were set to 100 percent. The percent change post-treatments were averaged and plotted for each oligomer size range indicated.

#### VP40 protein purification and Alexa-488 C_5_ Maleimide labeling

pET46-VP40-6xHis was transformed into Rosetta BL21 DE3 *E. coli* as previously described (2). The protein samples were purified following affinity chromatography using size exclusion chromatography on a HiLoad 16/600 Superdex 200 pg column (ÄKTA pure, Cytiva, Marlborough, MA). The desired fractions containing dimeric VP40 were collected. Protein concentration was determined with a BCA assay and protein was stored in 10 mM TRIS containing 300 mM NaCl, pH 8.0 for up to 14 days at 4°C. Labeling of VP40 cysteine residues was carried out using Alexa-594-C5-maleimide (See **doi:** https://doi.org/10.1101/2021.06.08.447555). VP40 dimer was treated with a 1.5-fold molar excess of Alexa-594-C5-maleimide dissolved in DMSO for 2 hours at room temperature in maleimide labelling buffer (20 mM NaPi solution, pH 7.4, containing 150 mM NaCl and 4 M Guanidine HCl). The labeling reaction was quenched by adding diothiothreitol (final concentration of 50 mM) and the labeled protein was separated from the non-conjugated dye on a HiLoad 16/600 Superdex 200 pg column. Fractions containing VP40 dimer were collected and concentrated. The labelling efficiency and protein concentration were estimated using a NanoDrop according to Invitrogen’s instructions.

#### Surface plasmon resonance assay

To determine the affinity of 6xHis-VP40 to LUVs with different PS species, SPR was performed. SPR experiments were performed at 25°C using a Biacore X100 as described previously (11). The apparent *K*_d_ of vesicle binding was determined using the non-linear least squares analysis:

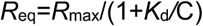

Where Req (measured in RU) is plotted against protein concentration (C). R_max_ is the theoretical maximum RU response and *K*_d_ is the apparent membrane affinity. Data were fit using the Kaleidagraph fit parameter of (m0*m1)/(m0+m2);m1=1100;m2=1. ΔRU data was normalized in GraphPad Prism 8 for windows (La Jolla, CA) and plotted in Kaleidagraph (Reading, PA).

#### Lipid binding assay

All lipids were purchased from Avanti Polar Lipids (Alabaster, AL). Lipid binding assays were performed as described previously (2). Briefly, lipid films were made by mixing chloroform soluble lipids and dried under nitrogen gas. In all experiments, lipid mixtures included 2% dansylPE. Dry lipid films were rehydrated with extrusion buffer (10 mM Tris, 160 mM NaCl, pH 8.0) containing 250 mM raffinose pentahydrate. Next, lipid vesicle solutions were extruded through a 0.2 μm membrane filter to form large unilamellar vesicles (LUVs) in solution. LUV solutions were diluted with LUV binding buffer (10 mM Tris, 160 mM NaCl) to reduce the raffinose to a final concentration of less than 16% (w/v%). 10 µM of protein were added to 2 mM LUVs, incubated for 30 minutes at room temperature, then pelleted at 75,000 x *g*, 30 minutes at 22°C. Supernatants (SN) were collected, and pellets (P) were resuspended in equal SN volumes and the samples were separated on SDS-PAGE. Protein band intensities on the SDS PAGE were analyzed with ImageJ for density in each sample. The percent protein in the pellet is represented as %bound (pellet/(pellet+SN)).

#### Liposome membrane fluidity assays

LUVs were prepared as described above and previously (12). In brief, lipid mixtures (see figure legend for more details) were made including the PA probe at 1:200 ratio (dye:lipid). Lipid films were rehydrated with extrusion buffer and a suspension of multilamellar vesicles were extruded through filters with a pore size of 0.2 μm (10 passages) and thereafter filters with a 0.1 μm pore size (10 passages). 0.6 mM of LUV were mixed with 1.5 μM 6xHis-VP40 (ratio 1:400, lipid:protein) for a final volume of 100 μl in a 96 well black/clear bottom plate. The fluorescence intensities were measured at time 0 minutes and at 30 minutes post-incubation of lipid:protein mixtures at 37°C, using Synergy Neo 2 (BioTek Instruments, Winooski, VT, USA). The dye was excited with λ_ex_= 430 nm and the fluorescence emission was collected from λ_em_= 450nm to λ_em_= 700 nm. The normalized intensities of three independent replicates were plotted for each conditions using GraphPad Software (San Diego, CA, USA).

#### EBOV infection of Huh-7 cells

All work involving EBOV was performed in the maximum containment laboratory (MCL) at the Rocky Mountain Laboratories (RML), Division of Intramural Research, National Institute of Allergy and Infectious Diseases, National Institutes of Health. Huh-7 cells were seeded in a 24-well plate the day before infection. Cells were infected with EBOV-GFP (13) at an MOI of 0.1 for 1 hour. The virus was then removed, and the cells were washed twice with DMEM. Fresh preparations of the drugs in calcium-free media were prepared and added to the cells. EBOV replication indicated by GFP was documented at 24-, 48-, and 72-hours post-infection. Work with EBOV-GFP was performed in the maximum containment laboratory at the Rocky Mountain Laboratories, NIAID, NIH following IBC-approved SOPs.

## Supplemental Figure Legends

**Figure S1.**
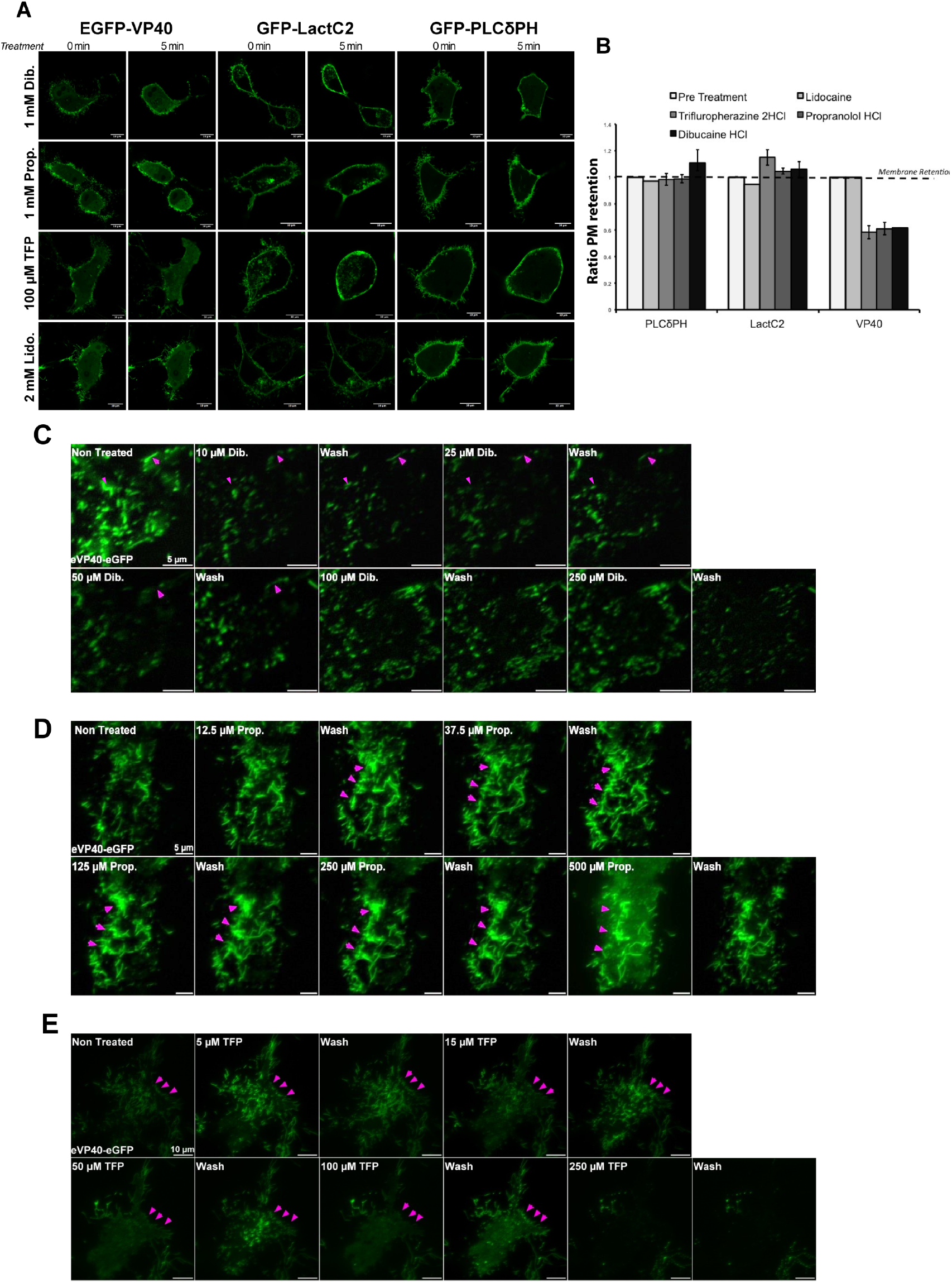
EBOV VP40 reversibly dissociates from the plasma membrane upon membrane fluidity increase despite the abundancy of PS and PI(4,5)P_2_. **A)** Representative images of GFP labelled VP40-WT, LactC2 and PLCδPH pre- and post-treatments at high drug concentrations. **B)** The ratio of plasma membrane retention of the different proteins from *A* quantified and normalized to pre-treatment conditions. The values are reported as mean ± SEM of three independent replicates. **C, D, E)** Representative TIRF imaging of the perfusion assay with dibucaine (C), propranolol (D) and trifluoperazine (E) where cells were treated with increasing concentrations of drugs for 10 min then washed with calcium free solution between treatments for 8 to 10 min to monitor the recovery of VP40 binding to the plasma membrane upon removal of drug treatment. Magenta arrows indicates areas of the plasma membrane were the protein recurrently bound and assembled after drug removal. Scale bar = 5 μm.

**Figure S2.**
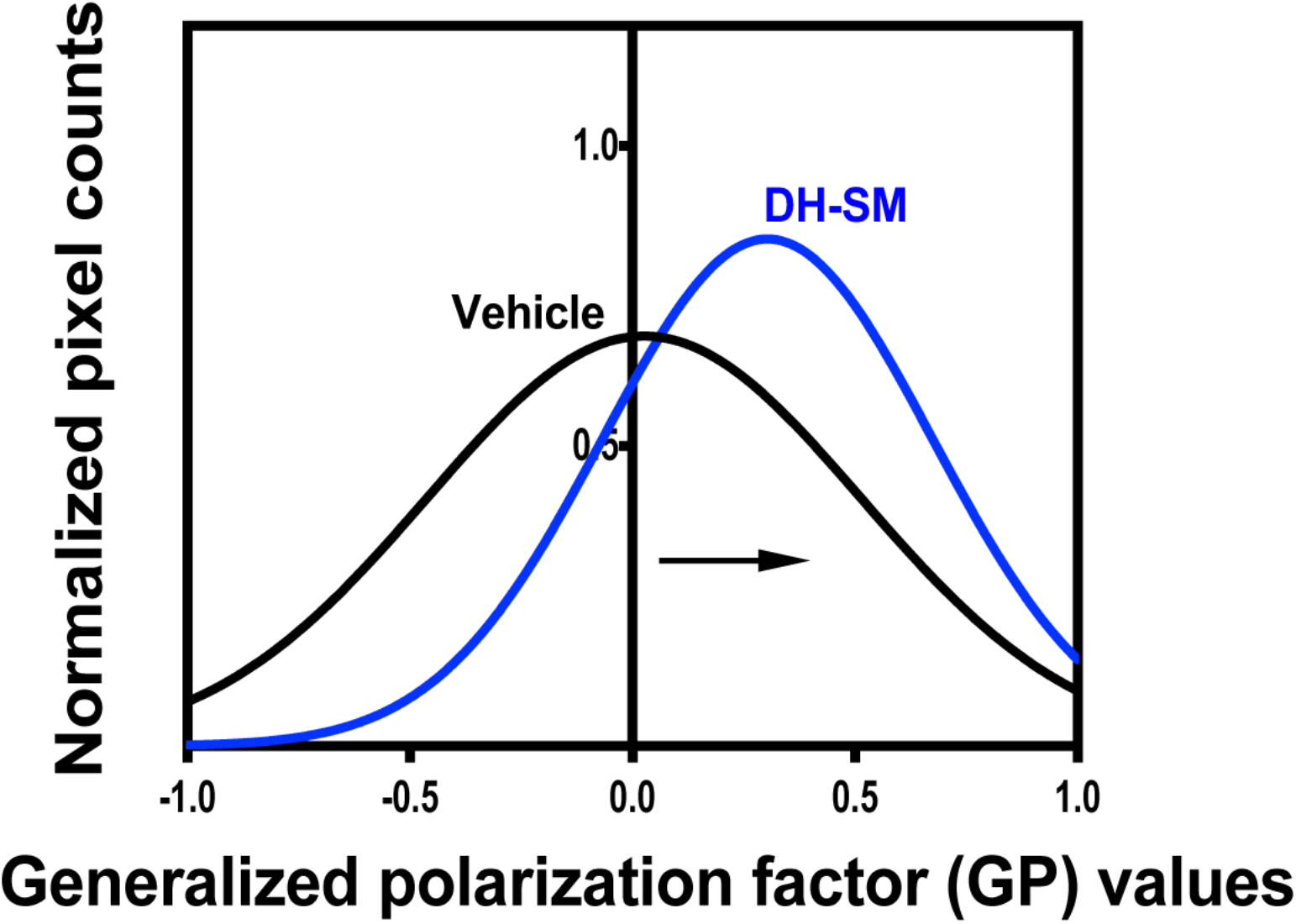
Experimental and fitted normalized generalized polarization (GP) distribution curves of Laurdan dye across the PM. HEK293 cells expressing EGFP-VP40 were treated with vehicle (black line) or 2.5 µM DH-SM (blue line).

**Figure S3.**
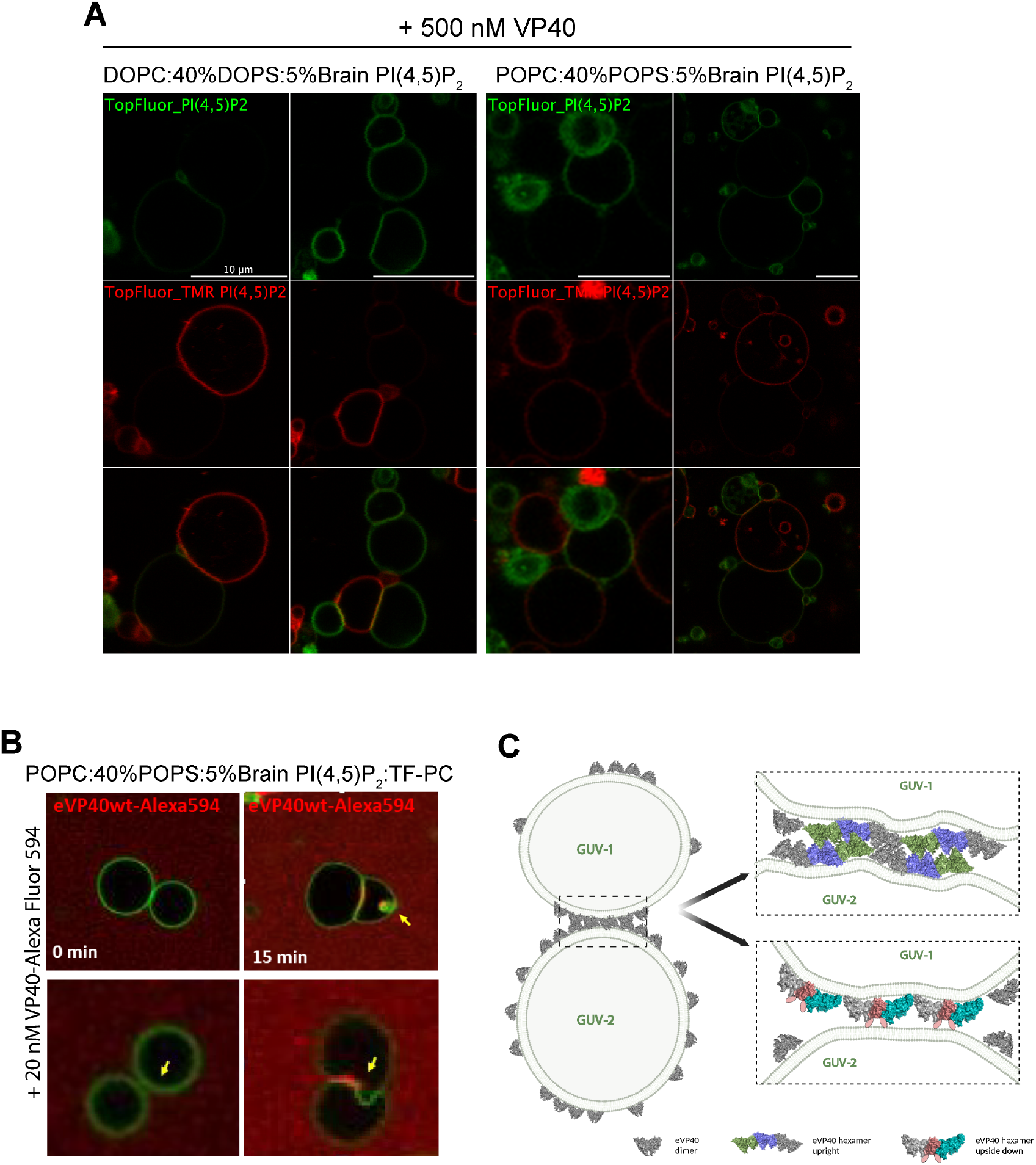
Recombinant EBOV VP40 induces membrane tethering and bending *in vitro*. A. Representative confocal images of a membrane fusion assay using GUV labelled with TopFluor PI(4,5)P_2_ (green) or TopFluor TMR-PI(4,5)P_2_ (red). The two GUV samples were mixed at equal molar ratio and incubated with 500 nM recombinant VP40 for 30 minutes to 1 hour at 37°C. B) Representative confocal imaging of membrane curvature, bending and vesiculation (white arrowheads) induced by recombinant VP40-AlexaFluor594 after 15 min incubation at 37°C. C) Caricatural illustration of potential VP40 assembly at the GUV membrane considering the two current models of EBOV matrix assembly.

**Figure S4.**
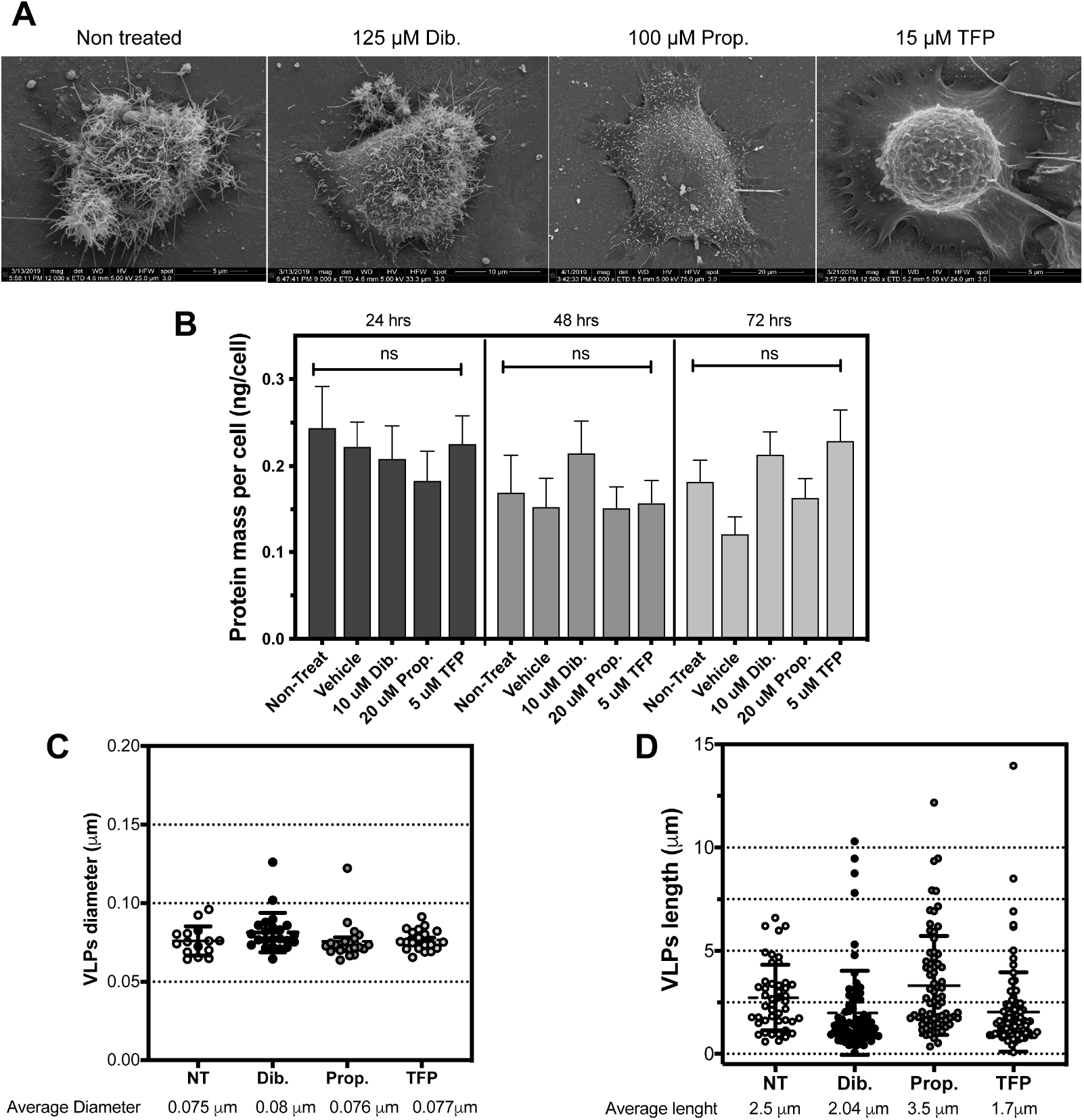
Cellular oligomerization defects of EBOV VP40 upon membrane fluidity increase reduces VLP budding. **A)** Representative SEM micrographs of HEK293 cells expressing EGFP-VP40 treated with the different FDA approved drugs. **B)** Average total protein mass per cell of HEK293 cells expressing EGFP-VP40 after 24-, 48- and 72-hours post-treatments with dibucaine, propranolol and trifluoperazine. The values are reported as mean ± SEM of three independent replicates. One-way ANOVA with multiple comparisons were performed (ns: non-significant). VLP diameter and length measurements were performed on TEM micrographs and are shown in **C)** and **D)**. The values are reported as mean ± SEM of three independent replicates. One-way ANOVA with multiple comparisons were performed.

**Figure S5.**
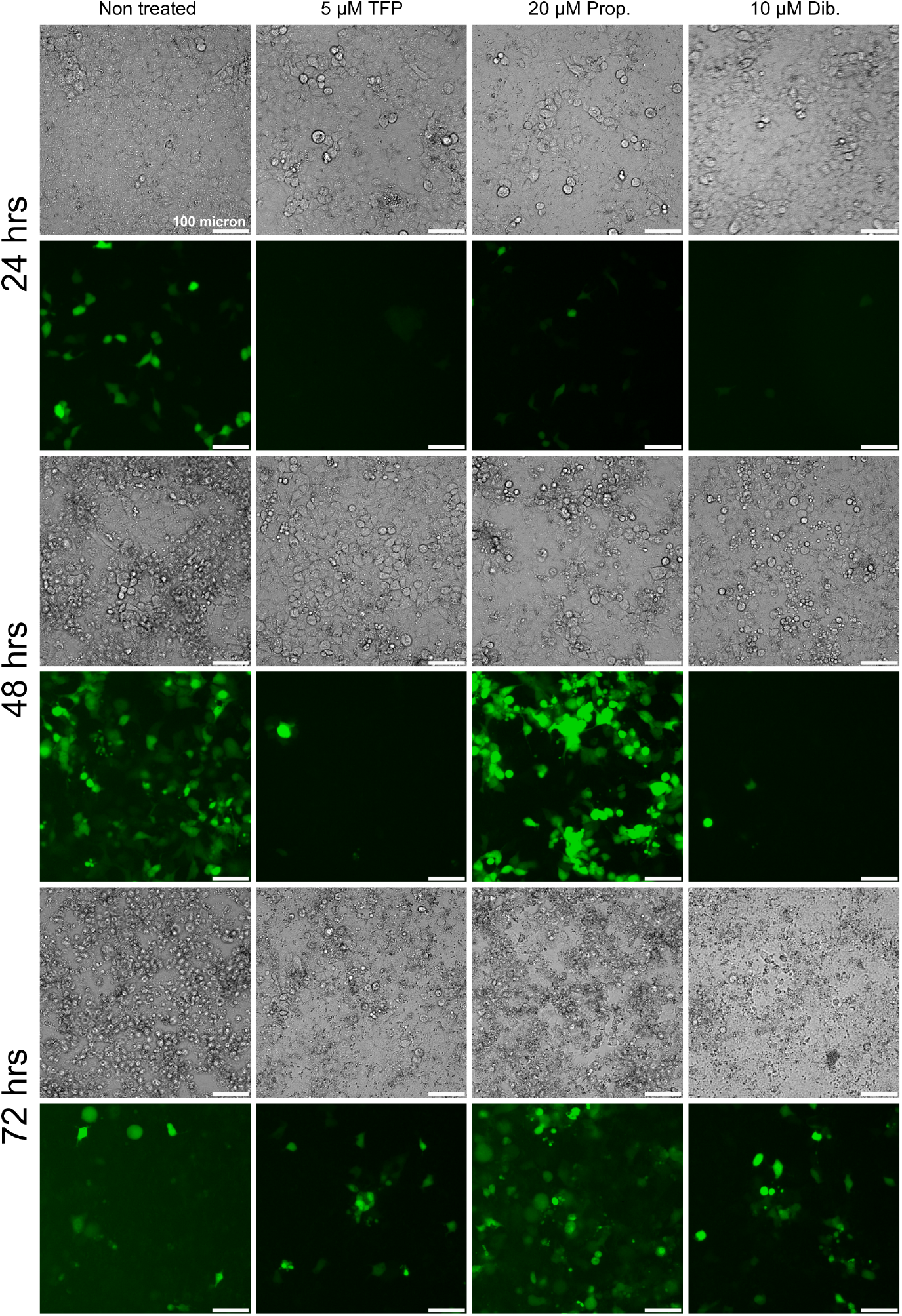
Effect of FDA-approved drugs on authentic EBOV infection and spread. Representative fluorescence images of Huh-7 cells infected with EBOV-GFP at the indicated MOI=0.1 and treated with the indicated concentration of different drugs to increase the plasma membrane fluidity. Cells were imaged at 24-, 48- and 72 hours post-treatments (green=EBOV; gray=cells). Scale bar = 100 μm.

